# High-resolution climate data reveals increased risk of Pierce’s Disease for grapevines worldwide

**DOI:** 10.1101/2024.03.06.583743

**Authors:** Àlex Giménez-Romero, Eduardo Moralejo, Manuel A. Matías

## Abstract

Range shifts in plant disease distributions are sensitive to scaling processes, but few crop case studies have included these predictions under climate change. High-quality wines are increasingly produced in topographically heterogeneous river valleys, whereby disease models that capture steep relief gradients become especially relevant. Here we show how non-linear epidemiological models more accurately reflect the threat of an emerging grapevine pathogen in areas with significant spatial gradients. By comparing the results of simulations using climate data with different spatial resolutions, we identify an increased risk of Pierce’s disease (PD), caused by the vector-borne bacterium *Xylella fastidiosa*, in wine regions globally. Over 100,000 vine presence records worldwide were analysed with respect to their closer risk-grid cell, observing an increase from 21.8% to 41.2% of the area at risk in European vineyards, from 5.6% to 47.2% in South Africa and to a lesser extent in other wine-growing regions. This general trend has been preceded by an accelerating rate of increase in risk within wine-growing areas. Our analysis demonstrates the importance of microclimatic conditions, highlighting previously unresolved risk zones in areas close to rivers and valleys, and the insufficiency of lower resolution data sets to capture complex climatic variations.

## Introduction

Climate plays a pivotal role in shaping the distribution and dynamics of agricultural pests and pathogens [1–5], with implications for global food security [6, 7]. As our climate undergoes unprecedented changes due to anthropogenic activities, agriculture faces multifaceted threats ranging from alterations in temperature and precipitation patterns to increased frequency of extreme weather events [8]. Such shifts create novel environments that may favour the proliferation of certain pests or pathogens while posing challenges to the survival of others [3, 9]. The consequences of these changes extend beyond immediate agricultural landscapes, reverberating through global food systems and posing significant challenges to the sustainability and resilience of food production [10].

Understanding the intricate relationships between climatic conditions, the pathosystem components, and the subsequent epidemiological dynamics is essential for developing effective strategies to mitigate and manage emerging agricultural challenges, especially in the face of changing environmental conditions. However, modelling disease epidemics is a complex task, as they are emergent phenomena resulting from non-linear interactions between disease components that also exhibit non-linear responses to changes in environmental variables [11–13]. Thus, while climate primarily determines the potential geographic range of each organism in the pathosystem, the development of epidemic outbreaks depends on favourable host-pathogen-vector-climate interactions that drive transmission chains.

It has long been recognised that ecological phenomena typically depend on the scale of description, particularly with regard to the effects of climate [14]. Climatic databases with finer spatial resolution are continuously being developed with the goal of allowing more accurate predictions [15]. Some recent studies have shown that the local climate experienced by individuals might deviate substantially from regional averages, with implications for the population dynamics of a forest herb [16]. Likewise, the choice of climate data affects the predictions of species distribution models (SDMs) [17]. In particular, the spatial resolution of the data can influence the predictions of invasion risk for some species [18]. It is therefore clear that the resolution of climate data will have a significant impact on predicting the risk of plant diseases and pests.

Among emerging pathogens *Xylella fastidiosa* (Xf) is considered one of the most dangerous phytopathogenic bacteria worldwide [19, 20]. It is naturally transmitted by xylem sap-feeding insects, such as sharpshooters and spittlebugs, and exhibits a broad host range that encompasses economically important crops such as grapevines, citrus, almonds and olive trees [20, 21]. The consequences of Xf diseases are devastating: about 200 million citrus trees are infected annually in Brazil [22], there are loses over $100 million annually in the grape industry in California [23] and approximately 21 million olive trees have been killed by the bacterium on the Apulia region in Italy [24]. Assuming massive spread throughout Europe, Xf has been projected to potentially contribute up to €5.2 billion of annual losses in the olive sector alone [25]. Overall, Xf diseases pose a major threat to agrosystems worldwide, highlighting the need for precise and predictive models to guide effective management practices.

Previous research has provided insights into the potential geographic range of Xf subspecies through SDMs [26, 27]. These models, however, have led to overestimates of risk by failing to account for the distribution and abundance of potential vectors necessary for disease transmission [28]. A quite different approach to mapping PD risk has been developed based on climate-driven epidemiological models with the option to integrate vector’s distribution information and the specificity of the Xf subsp. *fastidiosa* strain responsible for PD (hereafter Xf_PD_) [29]. This model correctly identifies areas in the United States with recurrent PD outbreaks and forecasts increasing epidemic risk in Mediterranean islands and coastlines with ongoing climate change.

Although risk maps based on hourly temperature data from the ERA5 have allowed fine adjustments in the calibration of the thermal response to Xf infection, these achievements have entailed losses in spatial resolution (0.1º spatial resolution) [29, 30]. Such limitation is particularly significant when dealing with vector-borne plant diseases like PD, where the interactions between the pathogen, vector, and host plants exhibit non-linear responses to climatic conditions. Subtle variations in temperature, humidity, or precipitation at the local scale thus can have profound effects on the reproduction and life cycles of the organisms involved and, hence, on the dynamics of disease transmission.

Topographical heterogeneity is a recognised issue in invasion biology, but has received little attention in crop science. Vineyards are increasingly located in valleys, ridges, hillsides and riverbanks usually with altitudinal and microclimatic gradients in short transects. They are therefore a remarkable example of a crop subject to scaling problems when studying ecological or epidemiological processes at regional and global scales. In this work, we address this spatial resolution limitation by modelling the risk of PD using high-resolution climate data from the CHELSA dataset [31]. The study period was deliberately chosen to include real data on temperature increases due to ongoing climate change. Our study shows a greater global risk of PD and a higher rate of risk increase, underscoring the urgency of reevaluating global strategies to prevent the spread of the pathogen with international trade in plant diseases.

## Results

### Global differences in PD risk between coarse and fine-grain climate data

We computed the risk of PD using the previously developed climate-driven epidemiological model [29] coupled with the CHELSA dataset [31], which features key climate variables (e.g. temperature and precipitation) at a high spatial resolution of 1 km and daily temporal resolution covering the period 1979-2016. The resulting spatial and temporal patterns of disease risk in the main wine-growing regions were compared with previous risk projections derived from the ERA5 dataset [32], characterised by an intermediate spatial resolution of 10 km and hourly temporal resolution [29]. Briefly, the model simulates the initial dynamics of the disease influenced by climatic variables and the presence of vectors, giving rise to a risk index, *r*, which represents the normalised growth rate of the infected population, where *r* = 1 is the maximum rate achieved at optimal climatic conditions (see Methods). Negative risk indices project an exponential decrease of the infected population (no risk), whereas positive values give rise to an outbreak, with higher values accounting for major incidence and potential severity. Risk categories emerge naturally from this formalism as No Risk (*r* ≤ −0.1) Transition (−0.1 < *r* ≤ 0.1), Low Risk (0.1 < *r* ≤ 0.33), Moderate Risk (0.33 < *r* ≤ 0.66) and High Risk (*r* > 0.66). Risk projections in Europe use the climatic suitability, *s*, of the main European vector, *P. spumarius* (see Methods), while for the rest of the world it is assumed that there are no risk-limiting effects due to the vector (*s* = 1), but only due to climatic conditions.

When contrasting model results derived from high- and medium-resolution data for the latest available time (2016), the disparity in risk projections extends beyond regional differences, showing a global increase in risk indices across wine-growing areas (Fig. 1 and Supplementary Fig. 1). Overall, these increases (Fig. 2) in the extension of PD risk areas ranged from 100,000 to 1 million km^2^ across viticulture regions worldwide. Transitions from no-risk to risk zones covered an area one order of magnitude larger than those in the opposite direction –from risk to no-risk (Fig. 2 and Table 1). In total, a surface of 4.6 million km-2 changed its risk category with the CHELSA database, representing about a 16% of the land area studied. In contrast, the largest decreases in the risk indices occurred mainly in the Southern Hemisphere, although with few exceptions most of these decreases remained within the risk zones (Fig. 1), while similar land expansions were observed to increase their risk category (low to moderate or moderate to high) (Fig. 2 and Table 1). The largest changes in risk indices occur in ecotones on both sides of the *r* = 0 line, as is clearly seen in the south-eastern United States, in coastal areas (e.g., southern Australia and northern California) due to higher resolution that better distinguishes between land and coast, and finally in the river valleys and slopes of mountain systems (Fig. 2 and Table 1).

**Table 1:** Changes in Pierce’s Disease Risk Zones in Different Viticulture Regions. The table illustrates transitions between risk and no-risk categories, as well as transitions among risk categories, highlighting the dynamic shifts in risk patterns across viticulture areas in Europe, the USA, South Africa, South America, and Australia. Risk increase refers to changes from low to moderate risk or from moderate to high risk. Likewise, risk decrease refers to changes from moderate to low risk or high to moderate risk.

|  | Risk to no-risk (km<sup>2</sup>) | Transition to risk (km<sup>2</sup>) | Risk decrease (km<sup>2</sup>) | Risk increase (km<sup>2</sup>) | No risk to risk (km<sup>2</sup>) | Total changes (km<sup>2</sup>) |
| --- | --- | --- | --- | --- | --- | --- |
| **Europe** | 1.91e+04 | 6.37e+04 | 3.58e+04 | 6.28e+04 | 2.04e+05 | 3.85e+05 |
| **USA** | 3.93e+04 | 1.37e+05 | 1.46e+05 | 2.37e+05 | 2.36e+05 | 7.95e+05 |
| **South Africa** | 2.26e+04 | 5.53e+04 | 3.56e+05 | 1.05e+05 | 2.77e+05 | 8.15e+05 |
| **South America** | 2.49e+05 | 3.79e+04 | 1.76e+05 | 1.50e+05 | 1.90e+05 | 8.03e+05 |
| **Australia** | 5.81e+04 | 8.49e+04 | 1.28e+06 | 1.55e+05 | 2.15e+05 | 1.79e+06 |

**Figure 1:**
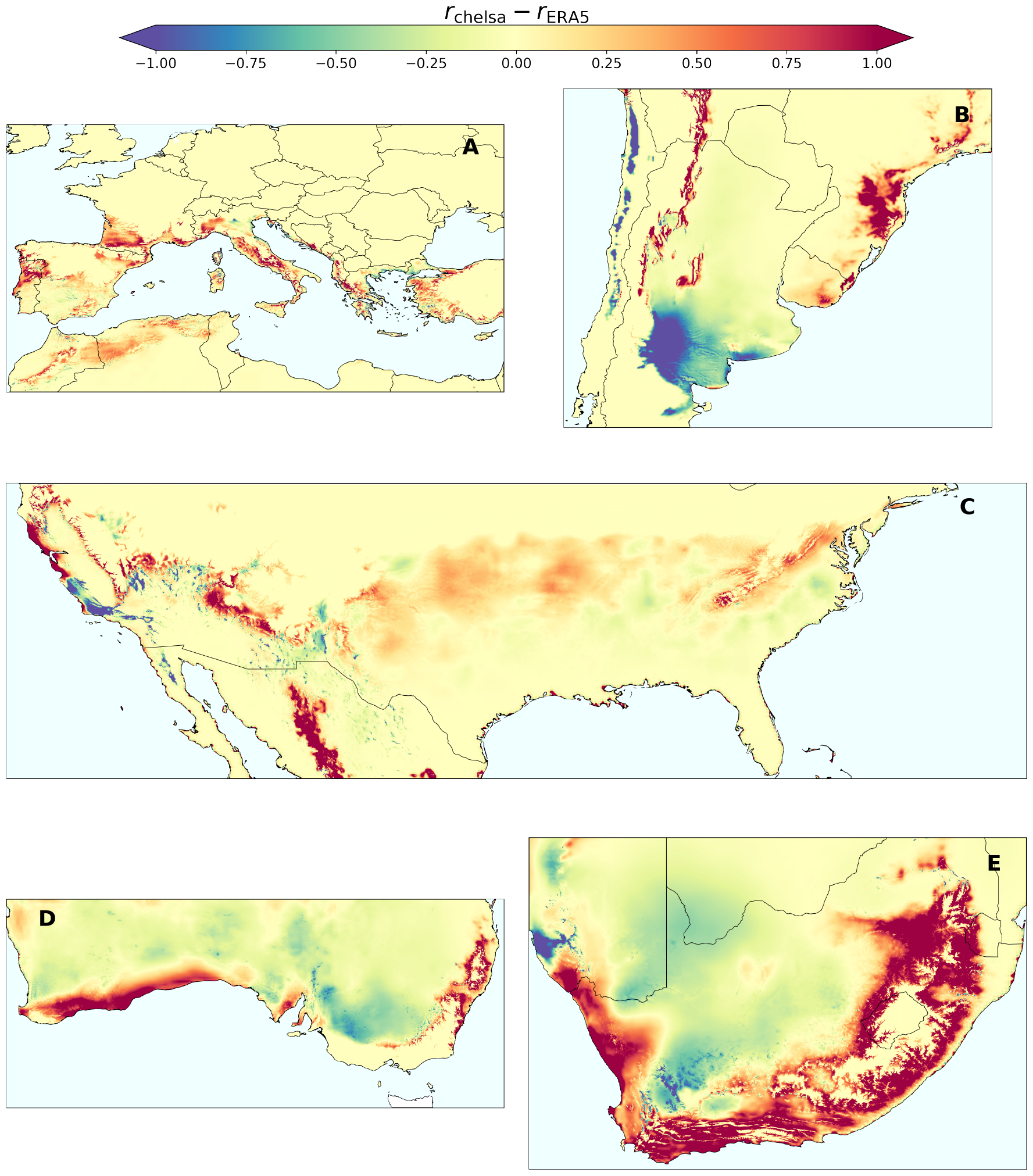
Difference in risk projections based on CHELSA (high-resolution, 1 km) and ERA5 (mid-resolution 10 km) datasets in global viticulture areas. (A) Europe (B) South America (C) United States (D) Australia (E) South Africa.

**Figure 2:**
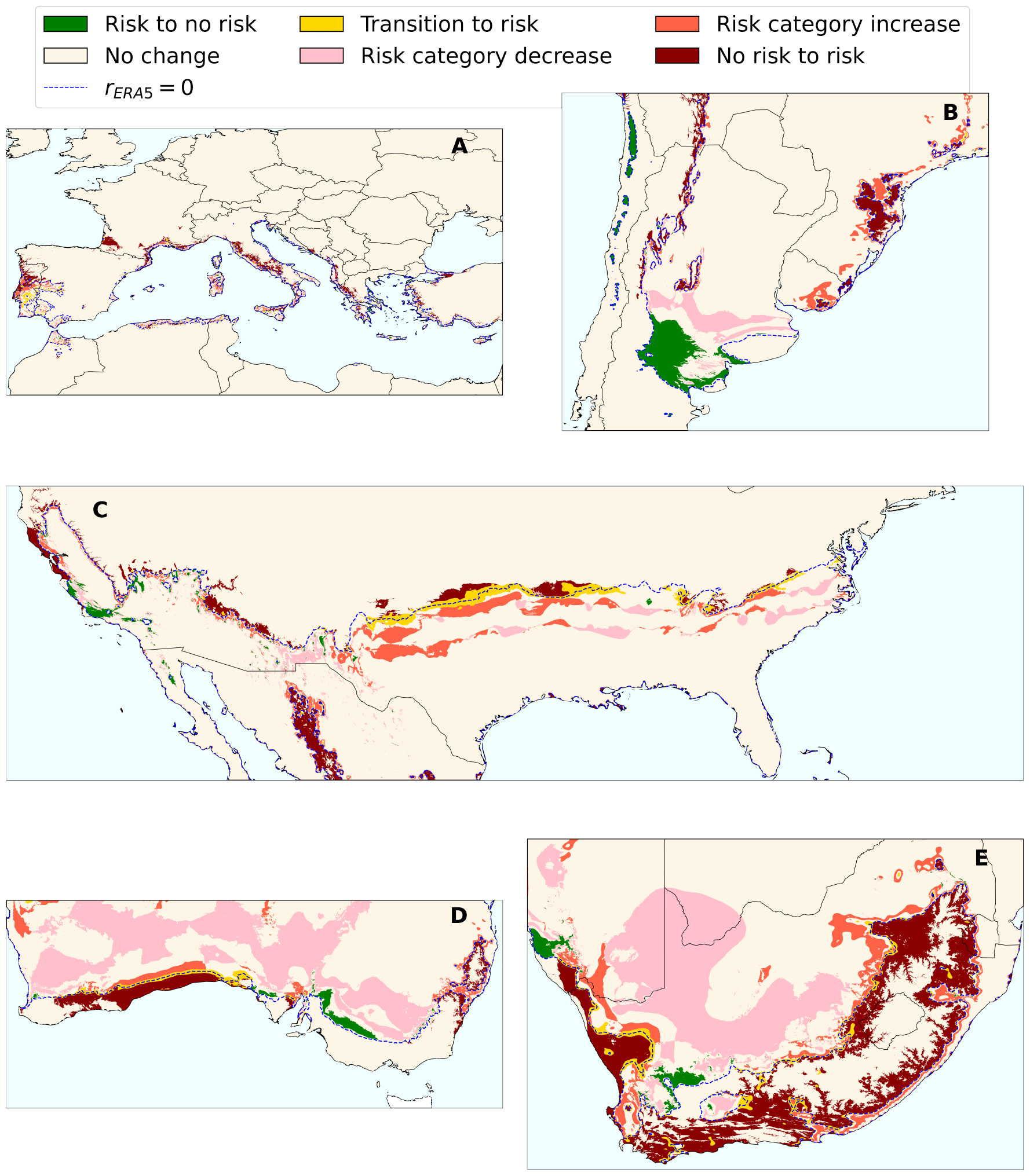
Changes in risk categories between CHELSA (high-resolution, 1 km) and ERA5 (mid-resolution 10 km) projections in global viticulture areas. (A) Europe (B) South America (C) United States (D) Australia (E) South Africa. Risk category increase refers to changes from low to moderate risk or from moderate to high risk. Likewise, risk category decrease refers to changes from moderate to low risk or high to moderate risk.

Next, we compared the temporal progression of the area at risk using both high and mid-resolution data over the entire available time span (1986-2016), considering that the risk for each year is computed based on the preceding seven years. We found a notable global surge in the rate increase of the area at risk within viticulture zones worldwide, practically doubling previous estimates (Supplementary Fig. 2). These results point to an accelerated pace at which the risk of PD is growing, compatible with the predictions of different global warming scenarios [33].

### Pierce’s Disease risk surges in previously unresolved microclimates

River valley vineyards are renowned for their high quality wines, such as Douro, Napa and Rhone and many others. It is therefore important to understand the risk of PD with climate change at a more detailed level. In our analysis, we have identified rivers and valleys as specific relief areas where a greater increase in PD risk is observed when employing CHELSA’s finer-scale climate data (Fig. 3). In some important wine-growing areas of southern Europe, we observed an abrupt emergence of risk zones previously classified as no-risk when using lower resolution climate data (Fig. 3). Such pronounced differences in risk patterns are highlighted for example in the fairly steep valleys and hillsides along the Douro River in Portugal, where the specific microclimatic conditions were previously obscured by the coarser resolution of the ERA5 data. These findings are particularly significant for PD, as vineyards are often located in close proximity to rivers or valleys and their surroundings, creating microclimates that attenuate cold winters (black dots in (Fig. 3)). A gradual increase in the climatic suitability for PD in some river basins may thus favour the spread of the pathogen from coastal to interior areas of the continents, allowing interconnection between areas that would otherwise remain isolated. Coastal areas close to cool water masses may also undergo an increase in risk when using higher resolutions data, as exemplified in California (Fig. 3 e,f).

**Figure 3:**
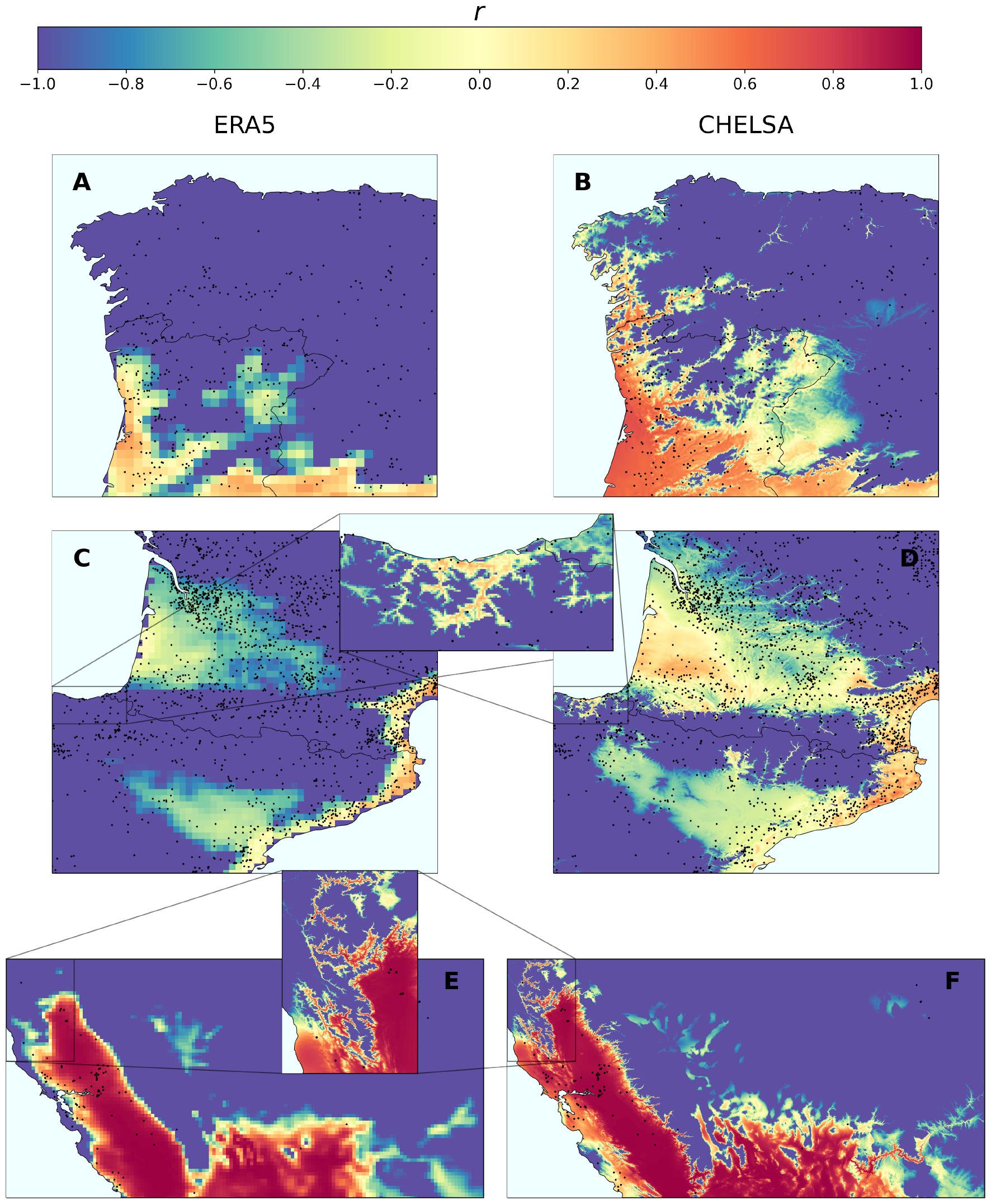
Effect of microclimatic conditions of rivers and valleys on Pierce’s Disease of grapevines. Comparison of the risk predicted using ERA5 mid-resolution dataset (A,C,E) and CHELSA high-resolution dataset (B,D,F). (A-B) North-western Iberian Peninsula. (C-D) Sourthern France and North-eastern Spain. (E-F) Western United States. Black dots represent grapevines (*Vitis vinifera*) presence data obtained from GBIF (see Methods).

Finally, to obtain a comprehensive assessment of the impact of microclimatic conditions on the risk of PD establishment, we collated a dataset of over 100,000 *Vitis vinifera* presence locations worldwide from GBIF [34], with a predominant concentration of points from Europe (Supplementary Fig. 3). Each data point was assigned a risk index based on the ERA5 and CHELSA projections, respectively, using the nearest pixel from each database. This approach revealed an increase in the risk indices associated with the vine locations (Fig. 4 A-D), mostly showing shifts towards higher risk indices (Fig. 4 A,E) from no risk to risk, or increases in risk category (low to moderate or moderate to high), while a negligible number of points decreased in risk category (Fig. 4 E). Such behaviour was common to all key viticulture regions studied, although the extent of increases differed between continents, with substantial expansion of vineyard areas at risk in Europe and South Africa (Table 2). Overall, our results emphasise the global relevance of microclimatic conditions in influencing the risk landscape for PD in viticultural areas (Table 2).

**Table 2:** Comparison of grapevine presence locations at risk in key viticulture regions using CHELSA and ERA5 datasets.

|  | N<sup>o</sup> points | risk CHELSA (%) | risk ERA5 (%) |
| --- | --- | --- | --- |
| **Europe** | 96102 | 41.2 | 21.8 |
| **USA** | 792 | 69.8 | 66.3 |
| **South Africa** | 36 | 47.2 | 5.6 |
| **South America** | 112 | 77.7 | 74.1 |
| **Australia** | 186 | 51.6 | 45.7 |

**Figure 4:**
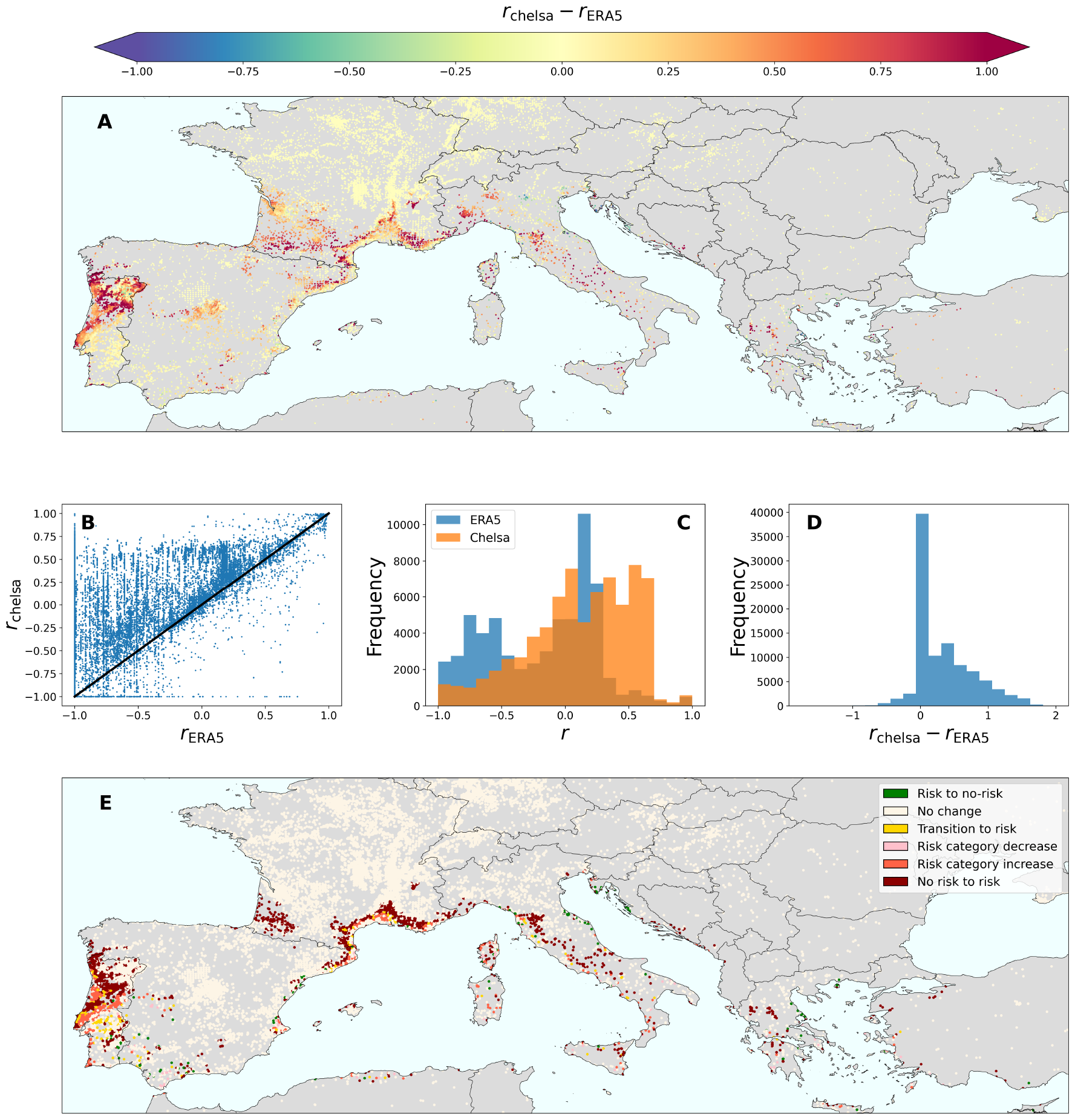
Impact of high-resolution climate data on the risk of Pierce’s Disease for grapevines worldwide. (A) Difference in risk indices in Europe, which accounts for the 96% of the points in the dataset. (B) Comparison of the risk indices derived from CHELSA and ERA5 datasets. Points with perfect agreement would lie in the solid black diagonal curve. (C) Histogram of risk indices derived from ERA5 (blue) and CHELSA (orange). (D) Histogram of the differences in risk indices between CHELSA and ERA5 datasets. (E) Changes in risk categories when using high-resolution climate data (CHELSA) with respect to mid-resolution data (ERA5). Risk category increase refers to changes from low to moderate risk or from moderate to high risk. Likewise, risk category decrease refers to changes from moderate to low risk or high to moderate risk.

## Discussion

Our study sheds light on the relevance of the spatial scale of observation in the intricate interplay between microclimatic conditions and the risk of PD for grapevines on a global scale. The use of high-resolution climate data reveals previously unrecognised local areas with microclimates conducive to the establishment of PD worldwide. Contrary to the simplistic assumption that higher resolution data might yield only marginal distinctions at regional levels, our study demonstrates that slight variations in climate data at local scales can lead to a global surge in disease risk. These increases not only affect the spatial distribution of risk, but also its temporal dimension, as suggested by the rate of increase in the surface area at risk. In the case of PD, we show that this rate nearly doubles when high-resolution climate data is considered compared to previous estimates obtained with mid-resolution data. Thus, our findings indicate a critical need for the use of local or high-resolution climate data in the assessment of disease risk, especially in areas characterised by diverse topography and even when only attempting to global estimates.

Such observed differences arise from the non-linear nature of disease dynamics and the response of the pathosystem components to environmental shifts [9, 11]. Therefore, models dependent on broader climate data may not capture the complexities of microclimates, resulting in an underestimation of disease risk. While this is not inherently negative, recognising these limitations helps to assume such risk estimates as a conservative lower bound until proven otherwise. Acknowledging these constraints is crucial for refining our understanding of disease dynamics and ensuring that our risk assessments are sufficiently cautious in the absence of more reliable data. Likewise, data coarsening procedures should be avoided, if possible, when modelling climate-driven disease dynamics, even in spite of computational efficiency. This recommendation applies not only to disease risk predictions but to all those in which non-linear functions depending on climate variables are present, such as species distribution models or phenological models [35].

Despite the valuable insights gained, our analysis heavily relies on the quality and resolution of the climate data from the CHELSA dataset [36]. While this dataset offers information at a high spatial resolution, the temporal dimension is limited to a daily frequency, which forces to apply an approximation to infer hourly data. Furthermore, the data may still be subject to biases or uncertainties inherent to the nature of the methodology employed in their construction. On the other hand, vector presence data is only accurately obtained for Europe, while an homogeneous presence is assumed in other viticulture areas. Additionally, the study primarily focuses on the effect of temperature conditions and the presence of potential vectors to determine the risk of Xf establishment, which may not encompass all possible contributing factors. Other variables, such as soil characteristics or vineyard management practices were not explicitly considered in this analysis, leaving room for additional complexities in the disease dynamics. Furthermore, the study predominantly examines the risk at a global scale, and the applicability of the findings to specific local contexts may vary.

Future research should aim to address the aforementioned limitations and provide a more comprehensive understanding of the multiple interactions influencing PD development in viticulture regions. Other factors influencing disease spread, such as human behaviour, land use changes, and ecological shifts, should also be explored, offering a more comprehensive and holistic view of the interplay between environmental conditions and disease vulnerability. The acceleration in the rate at which the risk of PD is growing calls for more research into control strategies to mitigate its impact on grapevine crops worldwide.

Although PD is currently restricted to North America and recently introduced in Taiwan [37], Mallorca (Balearic Islands, Spain) [38, 39] and Israel [40], since the mid-1990s climatic conditions are increasingly conducive to the establishment of PD in southern Europe [29]. For example, with the increase in the resolution of climate data our model predicts the recent detection of PD in Portugal [41], which was not anticipated using the ERA5 data [29]. In a short time, it is foreseeable that there will be more epidemic outbreaks in vineyards in southern Europe if the entry of infested plants is not controlled. This not necessarily have to be vines but can also include other plants such as almond trees or ornamental plants [42].

Overall, our study contributes to the growing body of knowledge on the impact of climate on agricultural pests and pathogens, emphasising the importance of considering microclimatic conditions for a more deep understanding of disease dynamics. Future research should focus on developing comprehensive models that integrate high-resolution climate data, considering both the global and local factors that influence disease dynamics. This holistic approach will enable a more accurate prediction of disease risk, allowing for the development of targeted management strategies and the enhancement of global food security.

## Methods

### Climate data

Climate data was downloaded from two datasets for our analysis: the ERA5 dataset [30, 32] and the CHELSA dataset [31, 36]. ERA5 offers mid-resolution climate data with a spatial resolution of 10 km and hourly temporal resolution, while CHELSA provides high-resolution data with a spatial resolution of 1 km and daily temporal resolution. Both datasets exhibit global coverage and encompass crucial climate variables, such as temperature and precipitation. For our simulations, we used the mean hourly temperature data from ERA5 dataset and the maximum and minimum daily temperature data from CHELSA dataset.

### Vector climatic suitability

Vector climatic suitability data was obtained from [27], in which a Generalised Additive Model (GAM) is employed to calibrate the relationship of *P. spumarius* global occurrence with moisture index and maximum temperatures during summer index estimated from 1979 to 2013 using the CHELSA dataset.

### Vineyard data

To assess the risk of Pierce’s Disease in locations where grapevines are present, we collected a comprehensive dataset of over 100,000 *Vitis vinifera* presence data records from the Global Biodiversity Information Facility (GBIF) [34, 43]. We note that while the dataset spans the globe, 96% of the points are located in Europe (Supplementary Fig. 3).

### Climate-driven epidemiological model

We used the model developed in [29], which describes the initial exponential rise (or decrease) of infected plants at the onset of an epidemic based on the spatial distribution of the vector and the bacterial growth and survival processes mediated by temperature. The density of vectors at a given cell controls the number of new plants that will be inoculated with the bacterium, while the local temperature mediates the growth and survival processes of the in-plant bacterial population, leading the initial inoculation to an infection or not. These temperature-driven growth and survival processes are described with the accumulation of two metrics denoted *Modified Growing Degree Days* (MGDD) and *Cold Degree Days* (CDD). The base function to compute the MGDD is proportional to the Xf temperature-dependent growth rate and is defined by

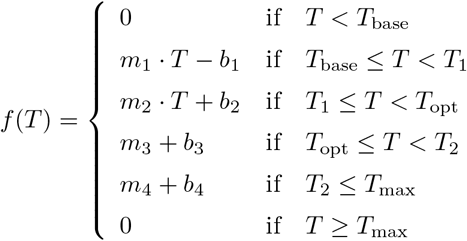

where *T*_base_ = 12 °C, *T*_1_ = 18, *T*_opt_ = 28 °C, *T*_2_ = 32 and *T*_max_ = 35 °C; the slopes are *m*_1_ = 0.66, *m*_2_ = 1, *m*_3_ = −1.25 and *m*_4_ = −3 and the intercepts are *b*_1_ = −8, *b*_2_ = −14, *b*_3_ = 4 and *b*_4_ = 105. MGDD are then computed as

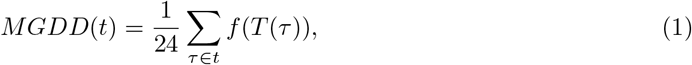

where *τ* is expressed in hours, *t* in years and we divide by 24 to obtain *MGDD*(*t*) in degree days. The accumulation period goes from the 1^st^ of April to the 31^st^ of October in the northern hemisphere and from the 1^st^ of November to the 31^st^ of March in the southern hemisphere.

CDD are computed between 1^st^ November and 31^st^ March in the northern hemisphere and between 1^st^ April and 31^st^ October in the southern hemisphere as

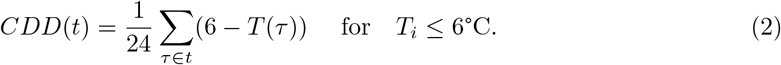

Altogether, the number of infected hosts is described by the following recurrence relation

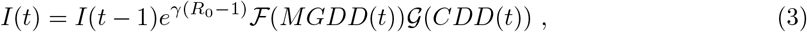

where *γ* is the death rate of infected vines, *R*_0_ is the basic reproduction number of the disease and ℱ(·) and 𝒢(·) are sigmoidal-like functions that relate the MGDD and CDD metrics to the probability of developing an infection from a given inoculation. Following [29], *R*_0_ in each cell *j* is related to the climatic suitability of the vector such that

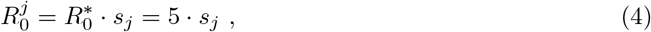

*γ* = 0.2 and the specific form of ℱ(·) and 𝒢(·) is given by

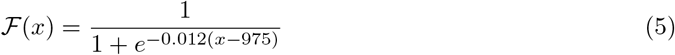

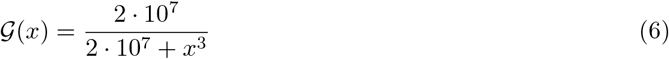

Finally, the risk index is derived as the effective growth rate of the infected population over the simulated time [29],

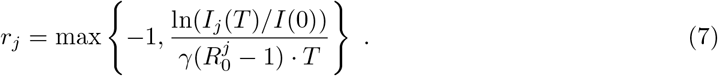

Because the typical time scale of the disease is 5 years (1*/γ*) [44], we simulate periods of 7 years. If more years are available to simulate, we perform a re-introduction of the disease as a single infected plant in each cell after each 7-year period [29].

The code used to run the model is freely accessible at GitHub [45].

### Model adaptation to daily temperature data

MGDD and CDD metrics were originally defined using hourly temperature data (Eqs. (1) and (2)) [29]. However, the CHELSA dataset only provide daily granularity. To overcome this limitation, we use a basic sinusoidal extrapolation relating maximum and minimum daily temperature to hourly temperatures,

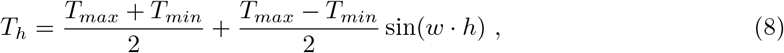

with *w* = 2*π/*24 and *h* ranging from 0 to 23. This approximation was validated in [33] with data from national meteorological stations in Spain (AEMET) using several locations and years, showing no differences between the use of hourly or daily temperatures to estimate MGDD and CDD. Similarly, the use of the approximation was validated across Europe using EURO-CORDEX data.

## Data availability

The MGDD, CDD and PD risk data generated in this study, both from ERA5 and CHELSA datasets, are available at [46]. Presence data on *Vitis vinifera* are available at GBIF [34].

## Code availability

The code for the climate-driven epidemiological model is available in a GitHub repository [45].

## Acknowledgements

AGR and MAM were supported through grant PID2021-123723OB-C22 (CYCLE) funded by MICIU/AEI/10.13039/501100011033 and by “ERDF A way of making Europe” and through grant CEX2021-001164-M (María de Maeztu Program for Units of Excellence in R&D) funded by MICIU/AEI/10.13039/501100011033.

We acknowledge the group of Alberto Fereres, especially his former post-doc Martin Godefroid, for providing the SDM-derived map of *Philaenus spumarius* from [28]

## Author contributions statement

AGR, EM and MAM conceptualised the project and conducted investigations; AGR processed the climate data; AGR performed the risk simulations and the analysis; AGR wrote the original draft; AGR, EM and MAM reviewed and edited the manuscript; EM and MAM supervised the project; MAM acquired funding.

## Competing interests

The authors declare no competing interests.

## Notes

https://zenodo.org/records/10579689

https://pdrisk.ifisc.uib-csic.es/future

### Competing Interest Statement

The authors have declared no competing interest.

