## Supplementary Information for "High-resolution climate data reveals increased risk of Pierce’s Disease for grapevines worldwide"

### Risk chelsa vs ERA5

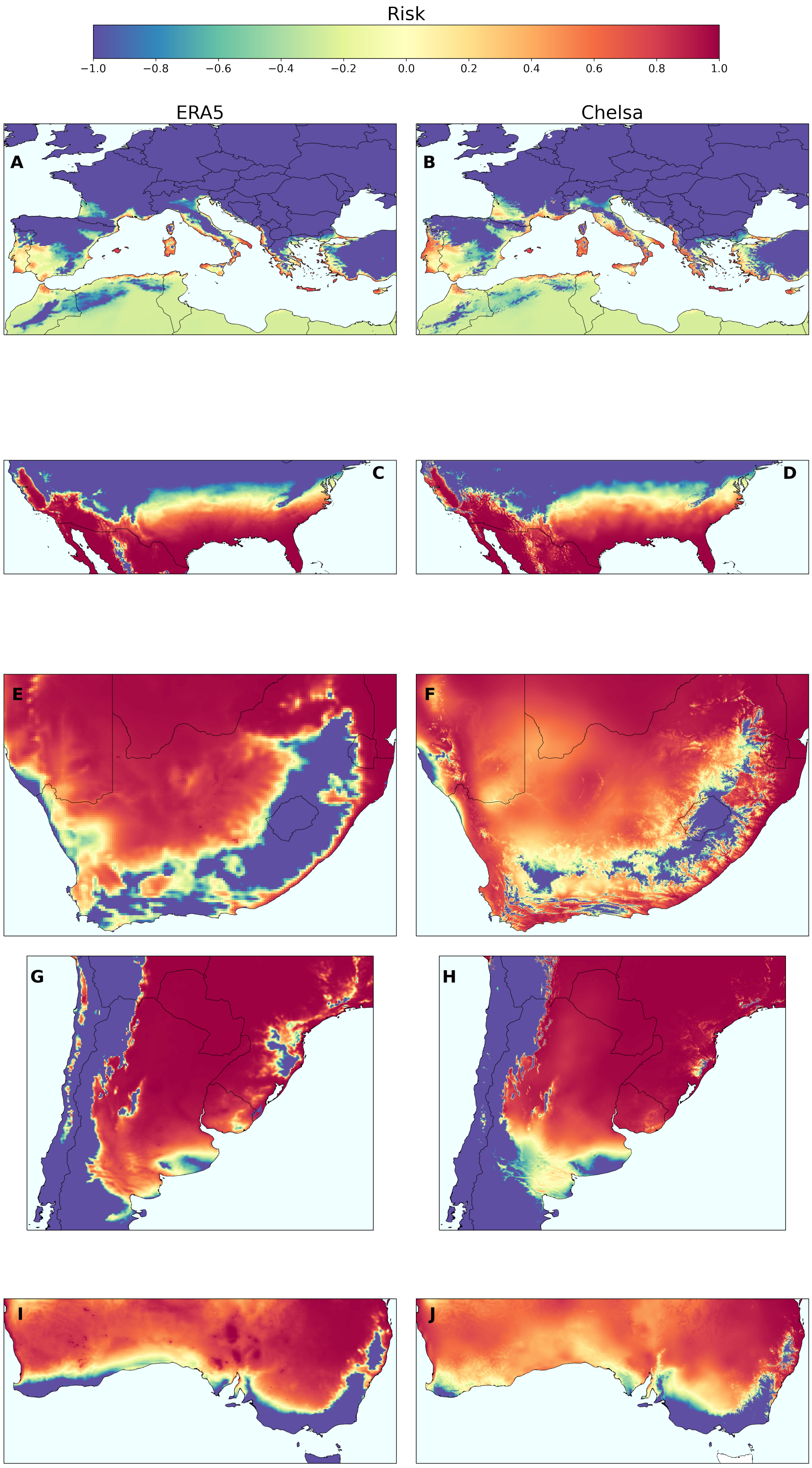


**Figure 1:** Comparison of risk indices obtained with ERA5 (mid-resolution – 10 km, left column) and CHELSA (high-resolution – 1 km, right column) datasets in Europe (A-B), United States (C-D), South Africa (E-F), South America (G-H) and Australia (I-J).

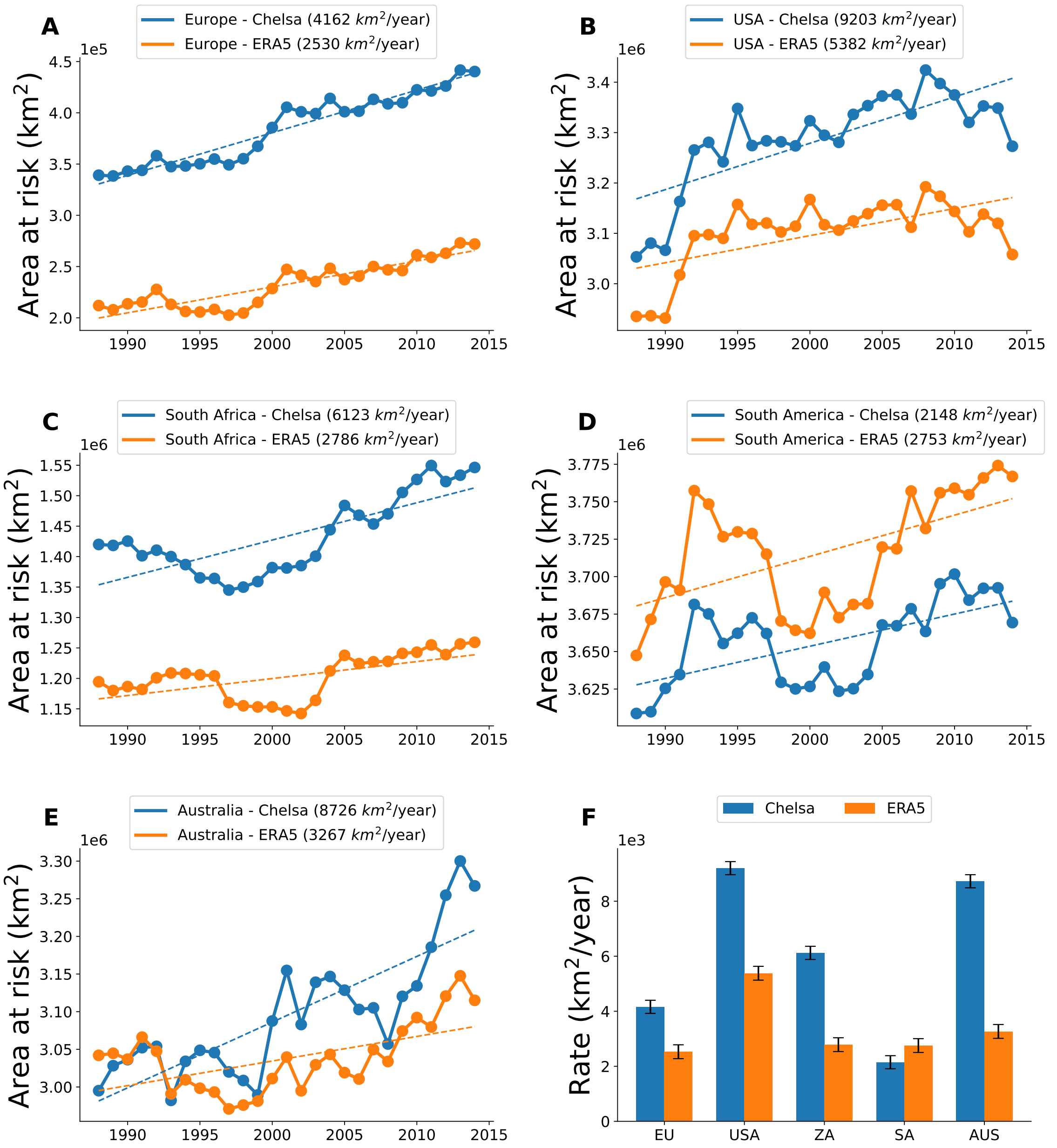


**Figure 2:** Difference in projected risk in increase rate based on CHELSA (high-resolution, 1 km) and ERA5 (mid-resolution, 10 km) datasets in global viticulture areas. (A) Europe (B) United States (C) South Africa (D) South America (E) Australia.

### *Vitis vinifera* global distribution

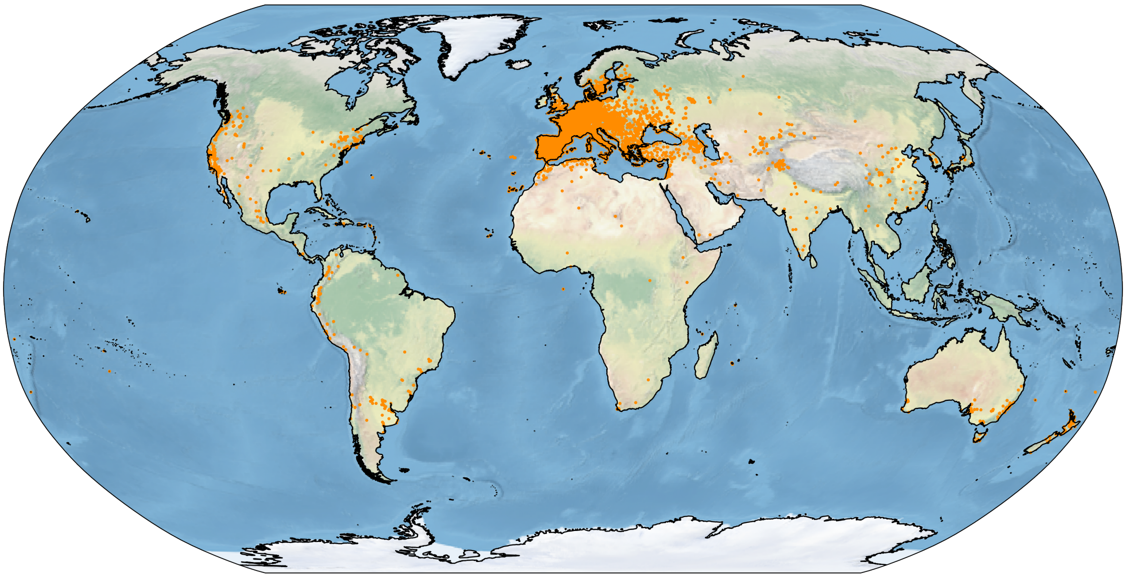

**Figure 3:** Presence locations of *Vitis vinifera* obtained from GBIF.
